# An update on eukaryotic viruses revived from ancient permafrost

**DOI:** 10.1101/2022.11.10.515937

**Authors:** Jean-Marie Alempic, Audrey Lartigue, Artemiy E Goncharov, Guido Grosse, Jens Strauss, Alexey N. Tikhonov, Alexander N. Fedorov, Olivier Poirot, Matthieu Legendre, Sébastien Santini, Chantal Abergel, Jean-Michel Claverie

## Abstract

One quarter of the Northern hemisphere is underlain by permanently frozen ground, referred to as permafrost. Due to climate warming, irreversibly thawing permafrost is releasing organic matter frozen for up to a million years, most of which decomposes into carbon dioxide and methane, further enhancing the greenhouse effect. Part of this organic matter also consists of revived cellular microbes (prokaryotes, unicellular eukaryotes) as well as viruses that remained dormant since prehistorical times. While the literature abounds on descriptions of the rich and diverse prokaryotic microbiomes found in permafrost, no additional report about “live” viruses have been published since the two original studies describing pithovirus (in 2014) and mollivirus (in 2015). This wrongly suggests that such occurrences are rare and that “zombie viruses” are not a public health threat. To restore an appreciation closer to reality, we report the preliminary characterizations of 13 new viruses isolated from 7 different ancient Siberian permafrost samples, 1 from the Lena river and 1 from Kamchatka cryosol. As expected from the host specificity imposed by our protocol, these viruses belong to 5 different clades infecting *Acanthamoeba spp*. but not previously revived from permafrost: Pandoravirus, Cedratvirus, Megavirus, and Pacmanvirus, in addition to a new Pithovirus strain.

## 1. Introduction

Ongoing international modeling and monitoring studies keep confirming that the continuous release of greenhouse gas (mostly CO_2_) due to human activities since the industrial revolution is causing significant climate change through global warming. It is now widely acknowledged that an average temperature increase of 1.5°C relative to 1850– 1900 would be exceeded during the 21st century, under all realistic circumstances [1] even though the adequacy of present climate models to predict regional changes remains in debate [2]. For instance, climate warming is particularly noticeable in the Arctic where average temperatures appear to increase more than twice as fast as in temperate regions [3]. One of the most visible consequences is the global thawing of permafrost at increasing depths [4, 5], the rapid erosion of permafrost bluffs [6,7], as well as erosion of deep and old permafrost by thaw slumping in hillslopes [8, 9]. This rapid permafrost thaw causes mobilization of ancient organic matter previously preserved for millennia in permafrost deep layers, a phenomenon most visible in Siberia where deep continuous permafrost underlays most of the North Eastern territories.

The thawing of permafrost has significant microbiological consequences. First, above freezing temperatures, the return of liquid water triggers the metabolic reactivation of numerous soil microorganisms (bacteria, archaea, protists, fungi)[10-14], exposing the organic material previously trapped in permafrost to decomposition, releasing additional CO_2_ and methane further contributing greenhouse gas to the atmosphere [5, 15, 16]. Yet, a more immediate public health concern is the physical release and reactivation of bacteria (or archaea) that have remained in cryptobiosis trapped in deep permafrost, isolated from the Earth’s surface for up to 2 million years [10, 17] (although a more consensual limit would be half a million years [18]). On a shorter time scale, the periodical return of anthrax epidemics devastating reindeer populations has been linked to the deeper thawing of the permafrost active layer at the soil surface during exceptionally hot summers, allowing century-old *Bacillus anthracis* spores from old animals burial grounds or carcasses to resurface [19-21]

One could imagine that very deep permafrost layers (i.e. million-year-old), such as those extracted by open-pit mining, could release totally unknown pathogens [22]. Finally, the abrupt thawing vertically operating along the whole wall of permafrost bluffs (consisting of specific ice-rich deposits called “yedoma”) such as seen in the Kolyma lowland or around the Yukon river Alaska, causes the simultaneous release of ancient microorganisms from frozen soils dating from the whole Holocene to the late Pleistocene (i.e. up to 120.000 years ago) [23]. Many culture-based and culture-independent studies (i.e. barcoding and/or metagenomics) have documented the presence of a large diversity of bacteria in ancient permafrost [10-12, 17, 24-28], a significant proportion of which are thought to be alive, although estimates vary greatly with the depth (age) and soil properties [17, 29, 30]. These bacterial populations include relatives of common contemporary pathogens (*Acinetobacter, Bacillus anthracis, Brucella, Campylobacter, Clostridia, Mycoplasma*, various *Enterobacteria, Mycobacteria, Streptococci, Staphylococci, Rickettsia*) [11, 12, 24, 29, 31]. Fortunately, we can reasonably hope that an epidemic caused by a revived prehistoric pathogenic bacterium could be quickly controlled by the modern antibiotics at our disposal, as they target cellular structures (e.g. ribosomes) and metabolic pathways (transcription, translation or cell wall synthesis) conserved during the evolution of all bacterial phyla [32], even though bacteria carrying antibiotic-resistance genes appear to be surprisingly prevalent in permafrost [26, 31, 33].

The situation would be much more disastrous in the case of plant, animal, or human diseases caused by the revival of an ancient unknown virus. As unfortunately well documented by recent (and ongoing) pandemics [34, 35], each new virus, even related to known families, almost always requires the development of highly specific medical responses, such as new antivirals or vaccines. There is no equivalent to “broad spectrum antibiotics” against viruses, because of the lack of universally conserved druggable processes across the different viral families [36, 37]. It is therefore legitimate to ponder the risk of ancient viral particles remaining infectious and getting back into circulation by the thawing of ancient permafrost layers. Focusing on eukaryote-infecting viruses should also be a priority, as bacteriophages are no direct threat to plants, animals, or humans, even though they might shape the microbial ecology of thawing permafrost [38].

Our review of the literature shows that very few studies have been published on this subject. To our knowledge, the first one was the isolation of Influenza RNA from one frozen biopsy of the lung of a victim buried in permafrost since 1918 [39] from which the complete coding sequence of the hemagglutinin gene was obtained. Another one was the detection of smallpox virus DNA in a 300-year-old Siberian mummy buried in permafrost [40]. Probably for safety/regulatory reasons, there was not follow up studies attempting to “revive” these viruses (fortunately). The first isolation of two fully infectious eukaryotic viruses from 30,000-y old permafrost was thus performed in our laboratory and published in 2014 and 2015 [41, 42]. A decisive advantage of our approach was to choose *Acanthamoeba spp*. as a host, to act as a specific bait to potentially infectious viruses, thus eliminating any risk for crops, animals or humans. However, no other isolation of a permafrost virus has been published since, which might suggest that these were lucky shots and that the abundance of viruses remaining infectious in permafrost is very low. This in fact is wrong, as numerous other *Acanthamoeba*-infecting viruses have been isolated in our laboratory, but not yet published pending their complete genome assembly, annotation, or detailed analysis. In the present article we provide an update on thirteen of them, most of which remain at a preliminary stage of characterization. These isolates will be available for collaborative studies upon formal request through a material transfer agreement. The ease with which these new viruses were isolated suggests that infectious particles of viruses specific to many other untested eukaryotic hosts (protozoans or animals) remain probably abundant in ancient permafrost.

## 2. Materials and Methods

### Permafrost sampling

The various on-site sampling protocols have been previously described in [31, 43] for samples #3 and #5 (collected in the spring 2015), in [13, 44] for sample #4, in [45] for sample #6, and [46, 47] for samples #7-9 (see Table 1).

**Table 1.** Samples and virus description

| Sample # | GPS coordinates | Description | Isolated virus |
| --- | --- | --- | --- |
| 1 | N 55°06'54"<br>E 159°57'48" | Surface soil, Shapina river<br>bank, Kamchatka<br>Modern | *Cedratvirus kamchatka* (strain P5) |
| 2 | N 62°06'23;92"<br>E 129°48'35" | Lena river, Yakutsk<br>Modern | *Cedratvirus lena* (strain DY0)<br>*Pandoravirus lena* (strain DY0) |
| 3 | N 61° 45' 39.1"<br>E 130° 28' 28.78" | Talik, -6.5 m below a lake,<br>Yukechi Alas [43]<br>Isolation: >53 y BP | *Pandoravirus talik* (strain Y4) |
| 4 | N 68°38'21.1"<br>E 159°3'20.67" | Melting ice wedge<br>Duvanny yar [23, 44]<br>Mixed ages | *Cedratvirus duvanny* (strain DY1)<br>*Pandoravirus duvanny* (strain DY1) |
| 5 | N 61° 45' 39.1"<br>E 130° 28' 28.78" | -16 m below a lake,<br>Yukechi Alas [43]<br>Isolation: >48,500 y BP | *Pandoravirus yedoma* (strain Y2) |
| 6 | N 74°13'00"<br>E 141°03'48" | Woolly mammoth stomach<br>content, Maly Lyakhovsky<br>Island [45]<br>Isolation: >28,600 y BP | *Pandoravirus mammoth* (strain Mm38) |
| 7 | N 70°43'25"<br>E 135°25'47" | Soil with mammoth wool<br>RHS paleolithic site,<br>Yana river left bank [46, 47]<br>Isolation: >27,000 BP | *Megavirus mammoth* (strain Yana14)<br>*Pithovirus mammoth* (strain Yana14)<br>*Pandoravirus mammoth* (strain Yana14) |
| 8* | N 70°43'25"<br>E 135°25'47" | Fossil wolf (*Canis lupus*)<br>intestinal content,<br>RHS paleolithic site [46, 47]<br>Isolation: >27,000 y BP | *Pandoravirus lupus* (strain Tums1) |
| 9* | N 70°43'25"<br>E 135°25'47" | Fossil wolf (*Canis lupus*)<br>intestinal content,<br>RHS paleolithic site [46, 47]<br>Isolation: >27,000 y BP | *Pacmanvirus lupus* (strain Tums2) |

\* Same location, but different frozen remains.

Liquid sample #2 and #4 were collected in pre-sterilized 50 ml Falcon tube in august 2019, as well as sample #1 consisting of surface soil without vegetation from the Shapina river bank collected on 07/15/2017 and since maintained frozen at - 20°C in the laboratory.

### Sample preparation for culturing

About 1g of sample is resuspended in 40 mM Tris pH 7.5, from 2-10% V/V depending on its nature (liquid, mud, solid soil) and vortexed at room temperature. After decanting for 10 minutes, the supernatant is taken up, then centrifugated at 10,000 g for one hour. The pellet is then resuspended in 40 mM Tris pH 7.5 with a cocktail of antibiotics (Ampicillin 100μg/mL, Chloramphenicol 34μg/mL, Kanamycin 20μg/mL). This preparation is then deposited one drop at a time onto two 15 cm-diameter Petri dishes (Sarsted 82.1184.500) one previously seeded with *Acanthamoeba castellanii* (Douglas) Neff (ATCC 30010TM) at 500 cells/cm^2^, the other with *A. castellanii* cells previously adapted to Fungizone (Amphotericin B, Gibco, Pasley, UK) by serial passages in presence of increasing concentration of the drug up to 2.5 μg/ml. Fungizone is used to inhibit the proliferation of viable microfungi known to be present in permafrost.

### Detection of virus infection

Changes in the usual appearance of *A. castellanii* cells (rounding up, non-adherent cells, encystment, change in vacuolization and/or refractivity) might eventually become visible after 72 h, but might be due to a variety of irrelevant causes such as overconfluency, the presence of a toxin, or the proliferation of bacteria or microfungi. Under a light microscope, the areas exhibiting the most visible changes are spotted using a p1000 pipetman. This 1 mL volume is then centrifugated (13,000 g for 30 minutes), the pellet resuspended in 100 μL and scrutinized under a light microscope. This sub-sample is also used to seed further T25 cell culture flasks of fresh *A. castellanii* cells.

### Preliminary identification of infecting viruses

Potential viral infections are suggested by intracellular changes (presence of cytoplasmic viral factories, nuclear deformation, lysis), or by the direct visualization of giant virus particles. Using a set of in-house-designed family-specific primers (Table 2), a PCR test is performed using the Terra PCR Direct Polymerase Mix (Takara Bio Europe SAS, Saint-Germain-en-Laye, France). Amplicons are then sequenced (Eurofins Genomics, Ebersberg, Germany) to confirm the presence of new isolates of a given acanthamoeba-infecting virus family (Table 2) suggested by their particle morphology and ultrastructural features.

**Table 2.**
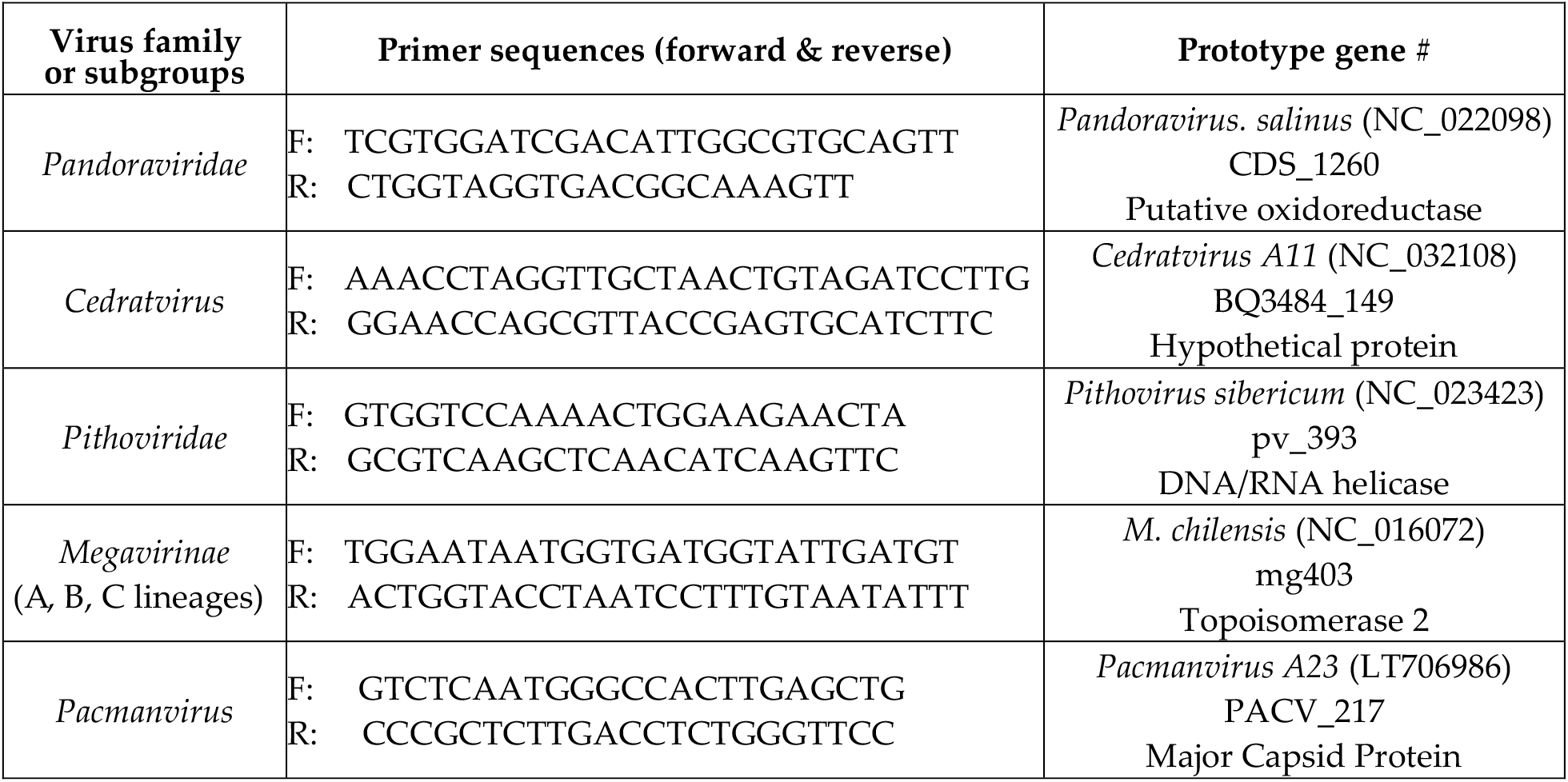
PCR primers used to identify the newly isolated viruses.

### Nomenclature of new isolates

We used the binomial format for the naming of virus species, where the genus name and a species epithet together form a unique species name. The genus name (e.g. “Pithovirus”) was attributed on the basis of concordant similarities with previously characterized amoeba-infecting viruses: genome sequences (PCR amplification using specific probes, partial or complete genome sequences), cell-infection patterns, and virion morphological features. The species epithet was chosen to reflect the location or nature of the source sample (e.g. “duvanny”). A strain name (e.g. “Tums1”) was added to further specify the precise sample (there might be several from the same location/source) from which the isolation was performed. Strain names can thus be shared by different species.

### Further characterization of new virus isolates

Positive subcultures are then reseeded and passaged in T25 then T75 cell culture flasks (Nunc™ EasYFlasks™, Thermofisher scientific, Waltham, MA USA) until the density/quantity of viral particles allows their further characterization by Transmission Electron Microscopy (TEM). New viral isolates of particular interest are then eventually cloned and their whole genome sequenced. The relationship of the new isolates to the other members of their cognate family was estimated using a phylogenetic clustering of the DNA-directed RNA polymerase largest subunit (RPB1) orthologous sequences. RPB1 is recognized as a convenient phylogenetic classifier for the nucleocytoplasmic large DNA viruses (phylum *Nucleocytoviricota*) [48].

### Viral genome sequencing

Virus cloning, virus particles purification using a cesium chloride gradient, and DNA extraction from approximately 5 × 10? purified particles (using the Purelink Genomic extraction mini kit, ThermoFisher) have been previously described [49]. Sequence data was generated from the Illumina HiSeq X platform provided by Novogene Europe (Cambridge, UK). Genome data assembly was performed in-house as previously described [49]. The draft genome sequences listed in Table 3-6 are provided as supplementary material (S1-S8).

**Table 3.** PCR identification of previous and new Pandoravirus isolates

| Virus | Accession # | Base pairs (contigs) | Amplicon identity* |
| --- | --- | --- | --- |
| *Pandoravirus salinus* (prototype) | NC_022098 | 2,473,870 | 100% (1203/1203) |
| *Pandoravirus celtis* | MK174290 | 2,028,440 | 93% (1128/1203) |
| *Pandoravirus quercus* | NC_037667 | 2,077,288 | 93% (1125/1203) |
| *Pandoravirus inopinatum* | NC_026440 | 2,243,109 | 93% (1133/1203) |
| *Pandoravirus dulcis* | NC_021858 | 1,908,524 | 92% (1122/1203) |
| *Pandoravirus neocaledonia* | NC_037666 | 2,003,191 | 86% (1045/1203) |
| *Pandoravirus macleodensis* | NC_037665 | 1,838,258 | 86% (1039/1203) |
| ***Pandoravirus lena* (strain DY0)** | - | 2,030,260 (6) | 93% (1131/1203) |
| ***Pandoravirus duvanny* (strain DY1)** | - | unassembled | 92% (1114/1203) |
| ***Pandoravirus talik* (strain Y4)** | - | 1,817,546 (1) | 92% (1114/1203) |
| ***Pandoravirus mammoth* (strain Yana14)** | - | not yet sequenced | 91% (1104/1203) |
| ***Pandoravirus yedoma* (strain Y2)** | - | not yet sequenced | 91% (1095/1203) |
| ***Pandoravirus lupus* (strain Tums1)** | - | not yet sequenced | 91% (1092/1203) |
| ***Pandoravirus mammoth* (strain Mm38)** | - | 1,776,082 (2) | 86% (1040/1203) |

\*Computed from the pairwise alignment of various amplicon nucleotide sequences with that of the reference sequence in *P. salinus*. New isolates are indicated in bold.

### Design of virus-specific PCR primers

Clusters of protein-coding genes common to all known members of a viral family or clade were identified using Orthofinder [50]. The protein sequence alignments of these clusters were converted into nucleotide alignments using Pal2nal [51]. Statistics on the multiple alignments where then computed using Alistat [52] and sorted using the “most unrelated pair criteria”. The corresponding alignments were thus visually inspected to select the variable regions flanked by strictly conserved sequences suitable as PCR primers. The primers and their genes of origin are listed in Table 2.

## 3. Results

### 3.1 Pandoraviruses

Seven of the 13 new virus isolates reported in the present article were found to be new members of the *Pandoraviridae* family. Observed under the light microscope during the early phase of the isolation process, their proliferation in acanthamoeba cultures generated inhomogeneous populations of large ovoid particles (up to 1 μm in length and 0.5 μm in diameter). As for known pandoraviruses, the infection of *A. castellanii* cells was initiated by the internalization of individual particles via phagocytic vacuoles. Eight to 10 hours after infection, the Acanthamoeba cells become rounded, lose their adherence, and new particles appear in the cytoplasm. The replicative cycle ends with the cells lysis releasing about a hundred particles each. Using TEM, the particles appeared enclosed in a 70-nm thick electrondense tegument with a lamellar structure parallel to the particle surface and interrupted by an ostiole-like apex (Figure 1.A). In complement of these morphological features unique to the *Pandoraviridae* [53], PCR tests were performed to confirm the identification of the new isolates using family-specific sets of primers (Table 2) and the amplicons sequenced to evaluate their genetic divergence with other members of the family (Table 3). All new isolates were found to be significantly distinct from each others and from contemporary strains, albeit within the range of divergence (93%-86% nucleotide identity) previously observed (Table 3). Among these new isolates, 4 originated from radiocarbon-dated ancient permafrost: *Pandoravius yedoma* (strain Y2) (>48,500 y BP), *P. mammoth* (strain Mm38) (> 28,600 y BP), *P. mammoth* (strain Yana14)(>27,000 y BP), and *P. lupus* (strain Tums1)(>27,000 y BP). One originated from an unfrozen permafrost layer: *P. talik* (strain Y4), one from a melted mixture of permafrost layers (*P. duvanny* (strain DY1), and one from the muddy bank fo the Lena river: *P. lena* (strain DY0). Draft genomic sequence of *P. lena, P. talik*, and *P. mammoth* (strain Mm38) are provided as supplementary files S1-S3. Their large sizes fall in the range of previously sequenced pandoraviruses (Table 3). A clustering of *P. duvanny, P. lena, P. talik, and P. mammoth* (strain Mm38) within the Pandoraviridae family is shown in Figure 2.

**Figure 1.**
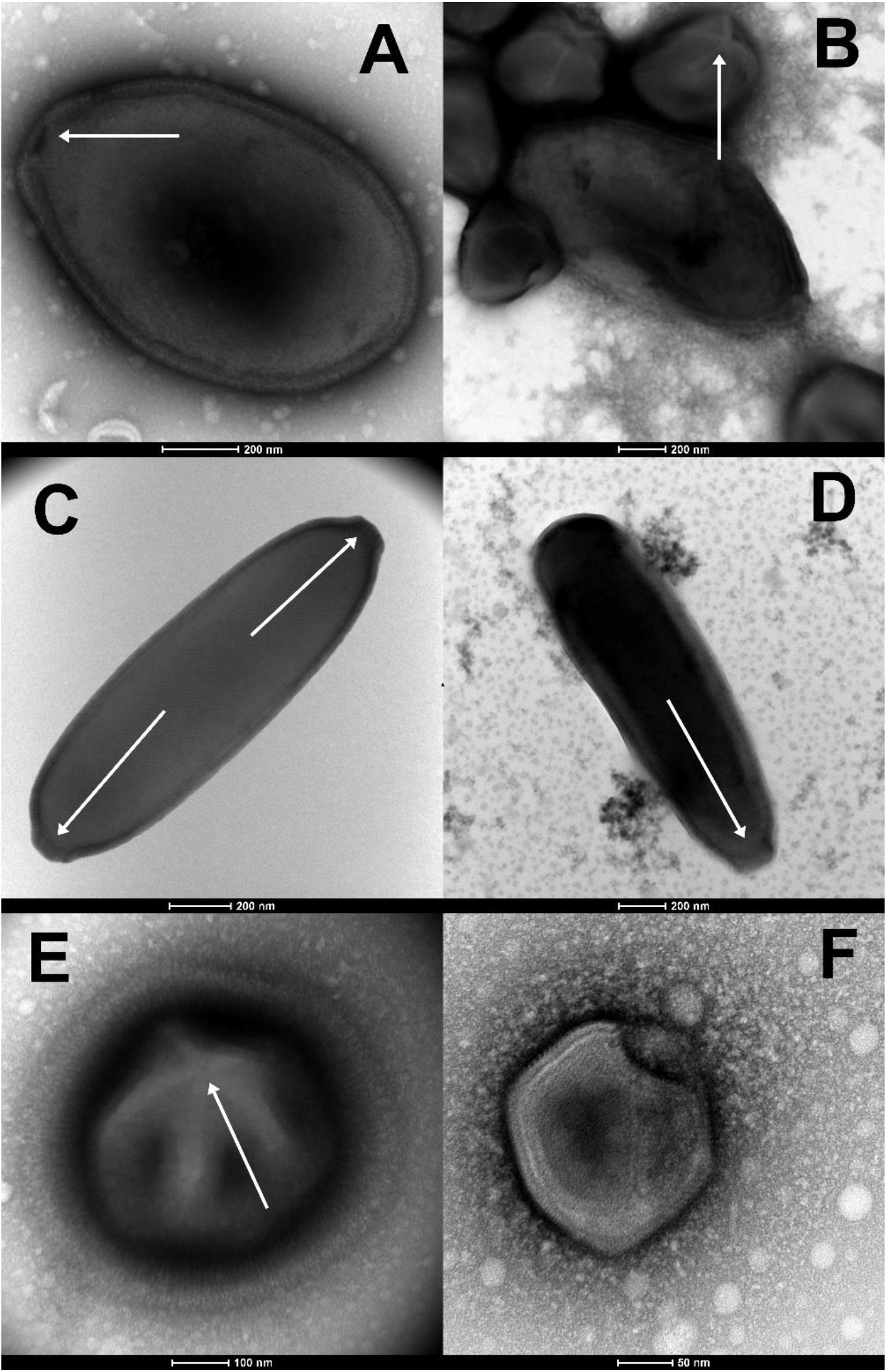
Morphological features guiding the preliminary identification of newly isolated viruses (negative staining, TEM). (**A**) The large ovoid particle (1,000 nm in length) of *Pandoravirus yedoma* (strain Y2) (sample #5 in Table 1) showing the apex ostiole (white arrowhead) and the thick tegument characteristic of the *Pandoraviridae* family. (**B**) A mixture of *Pandoravirus mammoth* (strain Yana14) oblate particles and of *Megavirus mammoth* (strain Yana14) icosahedral particles exhibiting a “stargate” (white starfish-like structure crowning a vertex, white arrowhead) as seen in sample #7 (Table 1). (**C**) The elongated particle of *Cedratvirus lena* (strain DY0)(1,500 nm in length) exhibits two apex cork-like structures (white arrowheads)(sample #2, Table 1). (**D**) The elongated particle of *Pithovirus mammoth* (1,800 nm in length) (sample 7, Table 1) exhibiting a single apex cork-like structure (white arrowhead). (**E**) The large (770 nm in diameter) “hairy” icosahedral particle of *Megavirus mammoth* (strain Yana14), showing the “stargate” (white arrowhead) characteristic of the *Megavirinae* subfamily (sample #7, Table 1). (**F**) The smaller icosahedral particle (200 nm in diameter) of *Pacmanvirus lupus* (strain Tums2) (sample #9, Table 1) typical of asfarviruses/pacmanviruses.

**Figure 2.**
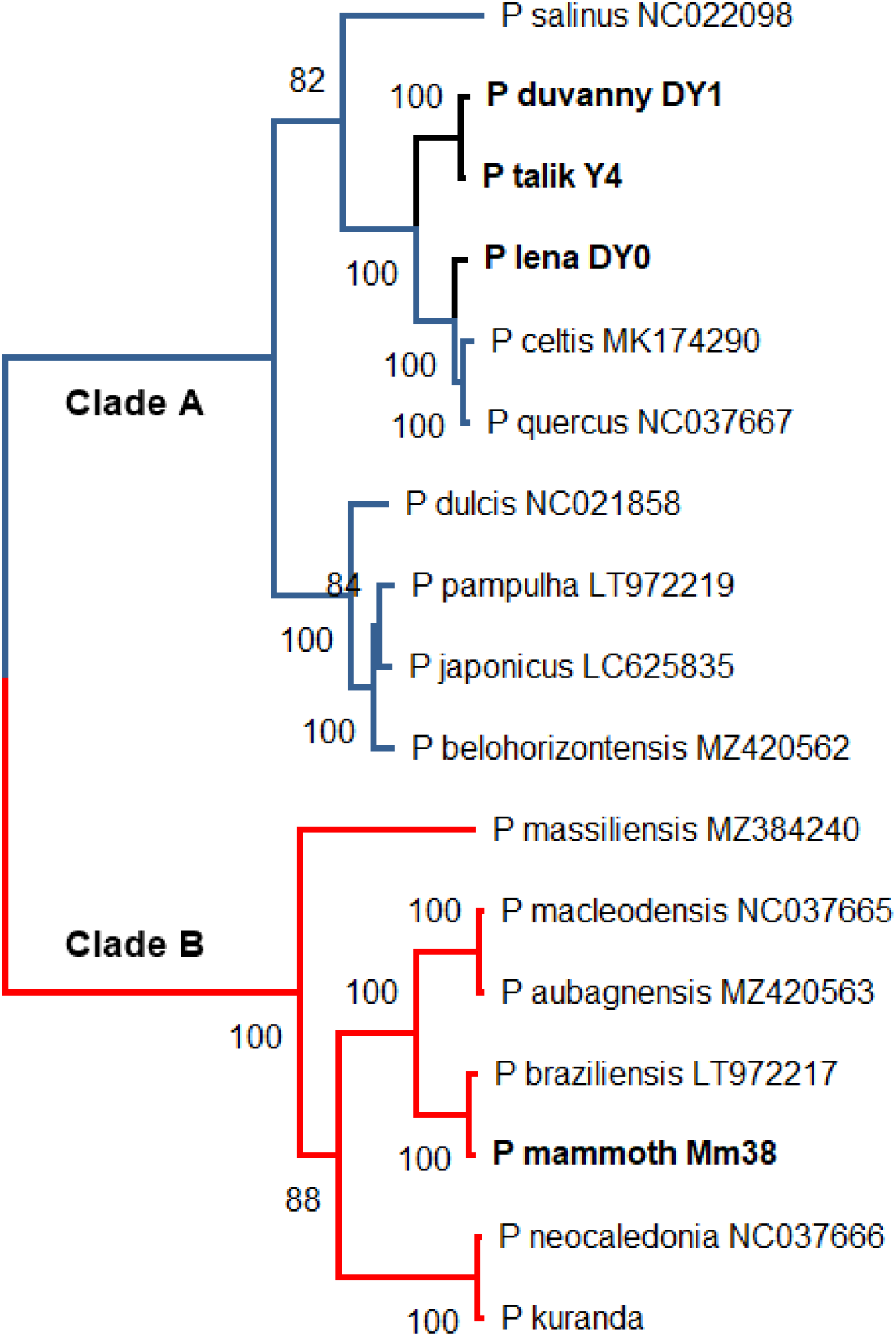
Maximum-likelihood phylogenetic relationships of the available Pandoravirus isolates. The tree (rooted at midpoint) was built using IQ-TREE (version 1.6.2) [54] from 2067 gap-free sites in the multiple alignment of 17 RNA polymerases (RPB1) protein (best fit model: “JTT+F+I+G4”). The permafrost isolates (in bold) are distributed between the two separate *Pandoraviridae* clades previously documented [**55**]. Accession numbers are indicated following the isolate name when available.

### 3.2 Cedratvirus and Pithovirus isolates

Three of the newly isolated viruses belong to the recently proposed “Cedratvirus” clade [56, 57] (a new genus or a new subfamily), within the *Pithoviridae* family [57]. One (*Cedratvirus lena* (strain DY0)) was cultivated from the same Lena river sample previously cited (Sample #2, Table 1), one (*Cedratvirus kamchatka* (strain P5)) from surface cryosol in Kamchatka collected during the summer (Sample #1, Table 1), and one (*Cedratvirus duvanny* (strain DY1)) from mud flowing into the Kolyma river at Duvanny yar, resulting from the thawing of permafrost layers of mixed ages (Sample #4, Table 1). One additional member of the *Pithoviridae* (*Pithovirus mammoth* (strain Yana14)) was isolated from a 27.000-y old permafrost sample containing a large amount of mammoth wool (Sample #7, Table 1). It is worth recalling that the prototype of this family was previously isolated from an ancient permafrost layer of more than 30,000-y BP [58]. Other members of this family are the most abundant in a recent metagenomic study of various Siberian permafrost samples focusing on eukaryotic viruses [59].

We recognized the new Cedratvirus and Pithovirus strains by their large ovoid particles, more elongated (up to 2 μm in length) than those of pandoravirus, with a much thinner wall, and their characteristic terminal cork-like structures (often two on each side for cedravirus particles) [56, 57](Figure 1.C, 1.D). Like previously described cedravirus/pithovirus, the new isolates enter the *Acanthamoeba* cells by phagocytosis. After ∼12 h of infection, mature viral particles were released by cell lysis. As previously noticed [58], the cell nucleus maintained its shape throughout the entire replication cycles.

In complement of these visual clues, PCR tests were performed to confirm the identification of the new isolates using two different clade-specific sets of primers (Table 2) and the amplicons sequenced to evaluate their genetic divergence with known members of the family (Table 4). All new isolates were found to be significantly distinct from each others and from contemporary strains, but within a range of divergence (94%-87%) consistent with that of previously characterized members of these clades (Table 4). In addition, we sequenced the genomes of the 3 new isolates. These draft sequences are provided as supplementary files S4-S6

**Table 4.** PCR identification of previous and newly Cedratvirus and Pithovirus isolates

| Virus | Accession # | Base pairs | Amplicon identity* |
| --- | --- | --- | --- |
| *Cedratvirus A11* (prototype) | NC_032108 | 589,068 | 100% (1239/1239) |
| *Cedratvirus lausannensis* | LT907979 | 575,161 | 94% (1176/1239) |
| *Cedratvirus kamchatka* | MN873693 | 466,767 | 88% (1091/1239) |
| ***Cedratvirus lena* (strain DY0)** | - | 465,544 | 87% (1090/1239) |
| ***Cedratvirus duvanny* (strain DY1)** | - | 472,117 | 87% (1087/1239) |
| ***Pithovirus sibericum* (P1084-T)** | NC_023423 | 610,033 | 100% (593/593) |
| ***Pithovirus mammoth* (strain Yana14)** | - | 610,309 | 97% (581/593) |

\*Computed from the pairwise alignment of various amplicon nucleotide sequences with that of the reference sequence in *Cedratvirus A11*. New isolates are in bold.

### 3.3 Megavirus mammoth

*Megavirus mammoth* (strain Yana14) is the first *Mimiviridae* family [60-62] member ever rescued from ancient permafrost. It was isolated from the highly productive sample (dated > 27.000-y BP) exhibiting fossil mammoth wool (sample #7, Table 1) together with two other viruses: *Pithovirus mammoth* (Yana14) and *Pandoravirus mammoth* (Yana14).

The particles of *M. mammoth* (strain Yana14) exhibit all the morphological features characteristic of a member of subfamily *Megavirinae*: a large icosahedral capsid of about 0.5 μm in diameter, surrounded by an external layer of dense fibrils (up to 125 nm thick) (Figure 1.E)[60, 61]. These features (icosahedral symmetry, large size, fibrils, and stargate) are unique to *Mimiviridae* members, making their identification straightforward and unambiguous [63].

Like previously described members of the *Megavirinae subfamily* [62], the *M. mammoth* particles enter host cells by phagocytosis. Six to eight hours p.i. infected cells start rounding and losing adherence. New particles are then produced in very large cytoplasmic viral factories, leaving the cell nucleus intact. New virions are then released in large quantities (burst size ≈ 500) through cell lysis.

In complement of the above unambiguous observations, a PCR tests was performed to confirm the identification of the new isolate using a *Megavirinae* specific sets of primers (Table 2) and the amplicon sequenced to evaluate its genetic divergence with known members of the family. *M. mammoth* was found to be a very close relative of the modern prototype *M. chilensis* (Table 5). Such very low levels of divergence are actually customary within the *Megavirus* genus (also referred to as the C-clade *Megavirinae*)[64]. A draft sequence of the *M. mammoth* (strain Yana14) genome is provided as supplementary file S7. A survey of this sequence shows that it encodes all the trademark proteins of the *Megavirus* genus [62]: the MutS-like DNA mismatch repair enzyme (ORF 570, 99% identical residues), the glutamine-dependent asparagine synthetase (ORF 434, 98% identical residues), and the 5 amino-acyl tRNA ligase: Ileu AARS (ORF 383, 99% ID), Asp AARS (ORF 771, 100% ID), Met AARS (ORF 798, 99% ID), Arg AARS (ORF 834, 99% ID), Cys AARS (ORF 837, 98% ID), Trp AARS (ORF 876, 96% ID), and Tyr AARS (ORF 944, 97% ID).

**Table 5.** PCR identification of *Megavirus mammoth* (strain Yana14) as a *Megavirinae* member

| Virus | Accession # | Base pairs | Amplicon identity* |
| --- | --- | --- | --- |
| *Megavirus chilensis* | NC_016072 | 1,259,197 | 100% (1497/1497) |
| *Megavirus vitis* | MG807319 | 1,242,360 | 99% (1493/1497) |
| ***Megavirus mammoth* (strain Yana14)** | - | 1,260,651 | 99% (1493/1497) |
| *Megavirus powai lake* | KU877344 | 1,208,707 | 93% (1396/1497) |
| *Megavirus baoshan* | MH046811 | 1,224,839 | 92% (1379/1497) |
| *Moumouvirus* | NC_020104 | 1,021,348 | 83% (1248/1497) |
| *Moumouvirus australiensis* | MG807320 | 1,098,002 | 82% (1244/1497) |
| *Mimivirus* | NC_014649 | 1,181,549 | 77% (1158/1497) |

\*Computed from the pairwise alignment of various amplicon nucleotide sequences with that of the reference sequence in *Megavirus chilensis*. The new isolate is in bold.

### 3.4 Pacmanvirus lupus

*Pacmanvirus* is a clade of recently discovered *Acanthamoeba*-infecting viruses distantly related to the African swine fever virus, until then the only known members of the *Asfarviridae* family that infects pigs [65]. We now report the isolation of a third member of this newly defined group from the frozen intestinal remains of a Siberian wolf (*Canis lupus*) preserved in a permafrost layer dated >27,000-y BP. At variance with the other truly giant viruses (i.e. exhibiting unusually large particles), their icosahedral virions are about 220 nm in diameter (Figure 1.F), hence not individually discernable under the light microscope (Nomarski optic). In absence of recognizable specific features, *Pacmanvirus lupus* (strain Tums2) was initially identified by PCR using a specific set of primers (Table 2) and a survey of its draft genomic sequence (provided as supplementary file S8).

*Pacmanvirus lupus* genome consists of a double-stranded DNA linear molecule of 407,705 bp, comparable in size to that of the previously studied members of this group (Table 6). However, out of its 506 predicted protein-coding genes, only 241 (47.6%) exhibit homologs in the two previously sequenced Pacmanvirus genomes, and 221 (43.7%) are ORFans. Thus, if *Pacmanvirus lupus* appears definitely closer to pacmanviruses than to any other known viruses, its evolutionary distance is larger than usually observed within a subfamily or a genus. This large discrepancy in global gene content is consistent with the low similarity observed between various core genes of *Pacmanvirus lupus* and their homologs in other pacmanviruses and closest relatives (Table 7). The asfarviruses appear even more distant with half the number of genes and half the genome size, perhaps calling for a little more caution before definitely classifying pacmanviruses within the *Asfarviridae* (Table 7) [65]. Based on a comparison of RPB1 orthologs, *P. lupus* appears unambiguously clustered with other known members of genus Pacmanvirus (hence justifying its name) (Figure 3).

**Table 6.** PCR Identification of *Pacmanvirus lupus* (strain Tums2) as a new member of genus *Pacmanvirus*.

| Virus | Accession # | Base pairs | Amplicon identity* |
| --- | --- | --- | --- |
| *Pacmanvirus A23* | NC_034383 | 395,405 | 100% (470/470) |
| *Pacmanvirus S19* | MZ440852 | 418,588 | 93% (439/470) |
| ***Pacmanvirus lupus* (strain Tums2)** | - | 407,705 | <85% (399/468)<br>+ two large insertions |

\*Computed from the pairwise alignment of various amplicon nucleotide sequences with that of the reference sequence in *Pacmanvirus A23*. The new isolate is in bold.

**Table 7.** Closest virus relatives of Pacmanvirus *lupus*

| *Pacmanvirus lupus*<br>Predicted protein | *Pacmanvirus A23*<br>NC_034383<br>%ID (aa) | *Pacmanvirus S19*<br>MZ440852<br>%ID (aa) | Faustovirus<br>KJ614390<br>%ID (aa) | Kaumoebavirus<br>NC_034249<br>%ID (aa) | Asfarviruses<br>NC_044958<br>%ID (aa) |
| --- | --- | --- | --- | --- | --- |
| RNA polymerase<br>(RPB1) ORF 302 | 79%<br>(1124/1415) | 80%<br>(1130/1415) | 49%<br>(707/1434) | 42%<br>(598/1429) | 41%<br>(596/1457) |
| RNA polymerase<br>(RPB2) ORF 33 | 85%<br>(1093/1289) | 85%<br>(1104/1301) | 55%<br>(681/1241) | 44%<br>(528/1211) | 43%<br>(526/1228) |
| DNA polymerase<br>(POLB) ORF 265 | 65%<br>(1036/1591) | 65%<br>(1032/1591) | 37%<br>(513/1385) | 27.5%<br>(344/1250) | 33%<br>(382/1163) |
| Genome size |  |  |  |  |  |
| 407,705 bp | 395,405 bp | 418,588 bp | 457-491 kb | 351-363 kb | 172-191 kb |

**Figure 3.**
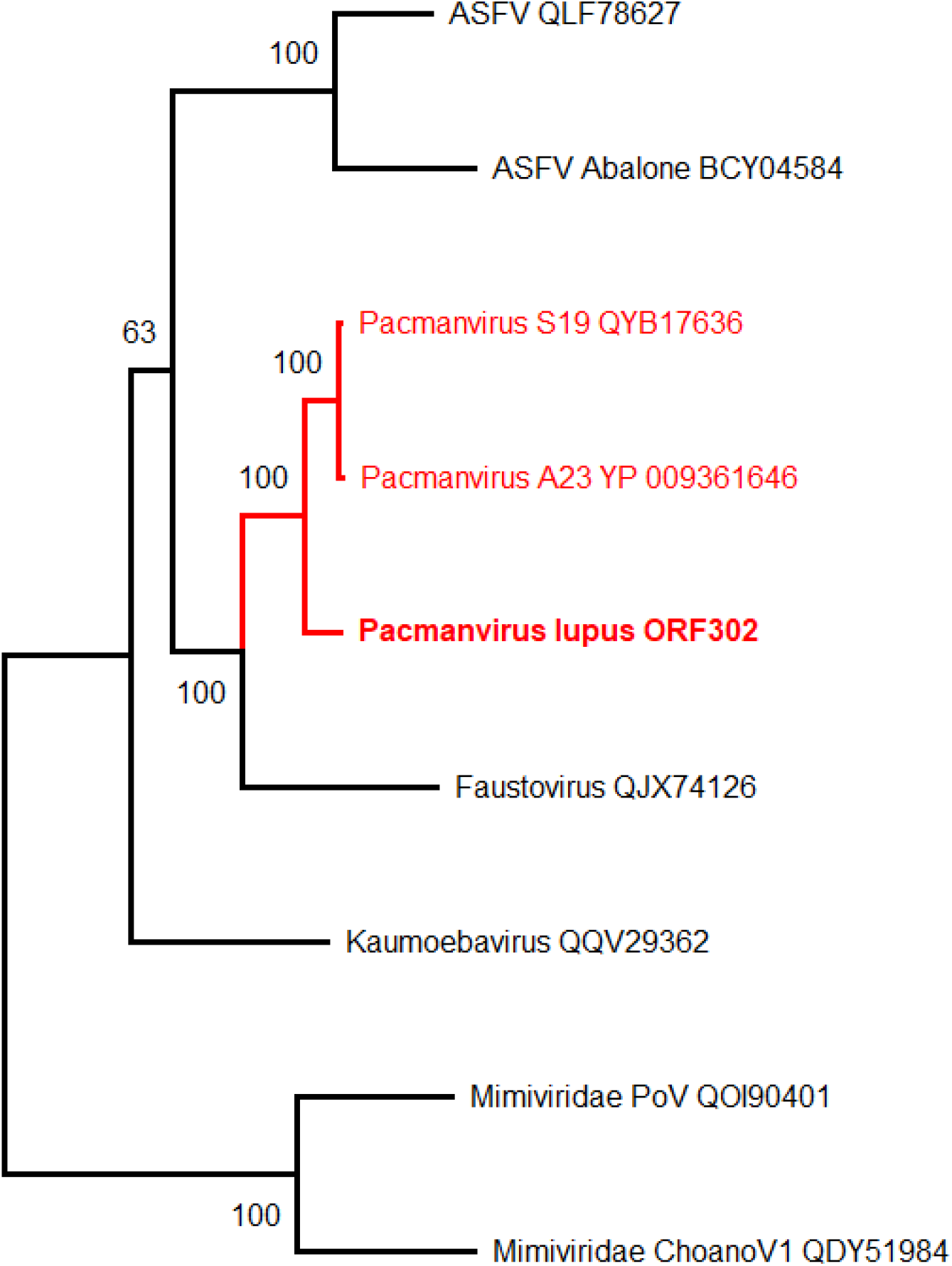
Maximum-likelihood phylogenetic relationships of the closest *Pacmanvirus lupus* relatives (using RPB1 homologs, Table 7). The tree (rooted at midpoint) was built using IQ-TREE (version 1.6.2) [54] (best fit model: “LG+F+I+G4”). The two closest *Mimiviridae* RPB1 sequences are used as an outgroup. The tree was built from 1314 gap-free sites in the multiple alignment of 9 RNA polymerases (RPB1) protein sequences. Although *Pacmanvirus lupus* is well clustered with other pacmanviruses, this clade (together with faustovirus) appears more as a sister group rather than *bona fide* members within the *Asfarviridae* (ASFV) family. Accession numbers are indicated following the isolate name when available

## 4. Discussion

Following initial reports published more than 5 years ago [41, 42], this study confirms the capacity of large DNA viruses infecting *Acanthamoeba* to remain infectious after more than 48,500 years spent in deep permafrost. Moreover, our results extend our previous findings to 3 additional virus families or groups: 4 new members of the *Pandoraviridae*, one member of the *Mimiviridae*, and one pacmanvirus (Table 1). One additional pithovirus was also revived from a particularly productive sample dated 27,000-y BP (sample#7, Table 1) exhibiting mammoth wool. Given these viruses’ diversity both in their particle structure and replication mode, one can reasonably infer that many other eukaryotic viruses infecting a variety of hosts much beyond *Acanthamoeba spp*. may also remain infectious in similar conditions. Genomic traces of such viruses were detected in a recent large-scale metagenomic study of ancient permafrost [59] as well as in Arctic lake sediments [66]. They include well documented human and vertebrate pathogens such as poxviruses, herpesviruses, and asfarviruses, although in lower proportions than protozoan infecting viruses.

In our recent metagenomic study [59], pandoraviruses are notably absent while they constitute the large majority of the viruses revived from permafrost and cryosols. Such a discrepancy might originate from the fact that the extraction of genomic DNA from their sturdy particles requires a much harsher treatment than for most other viruses. Their abundance in environmental viromes might thus be much larger than the small fraction they contribute to the DNA pool. Such DNA extraction bias may apply to many other microbes, and is a serious limitation to the validity of metagenomic approaches for quantitative population studies.

The types of viruses revived in our study are indeed the results of even stronger biases. First, the only viruses we can expect to detect are those infecting species of *Acanthamoeba*. Second, because we rely on “sick” amoeba to point out potentially virus-replicating cultures, we strongly limit ourselves to the detection of lytic viruses. Third, we favor the identification of “giant” viruses, given the important role given to light microscopy in the early detection of positive viral cultures. It is thus likely that many small, non-lytic viruses do escape our scrutiny, as well as those infecting many other protozoa that can survive in ancient permafrost [10].

However, we believe that the use of *Acanthamoeba* cells as a virus bait is nevertheless a good choice for several reasons. First, *Acanthamoeba* spp. are free-living amoebae that are ubiquitous in natural environments, such as soils and fresh, brackish, and marine waters, but are also in dust particles, pools, water taps, sink drains, flowerpots, aquariums, sewage, as well as medical settings hydrotherapy baths, dental irrigation equipment, humidifiers, cooling systems, ventilators, and intensive care units [67]. The detection of their virus may thus provide a useful test for the presence of any other live viruses in a given setting. Second, if many *Acanthamoeba* species can be conveniently propagated in axenic culture conditions, they remain “self-cleaning” thanks to phagocytosis, and are capable of tolerating heavy contamination by bacteria (that they eat) as well as high doses of antibiotics and antifungals. The third, but not the least, advantage is that of biological security. When we use *Acanthamoeba spp*. cultures to investigate the presence of infectious unknown viruses in prehistorical permafrost (in particular from paleontological sites, such as RHS [46, 47]), we are using its billion years of evolutionary distance with human and other mammals as the best possible protection against an accidental infection of laboratory workers or the spread of a dreadful virus once infecting Pleistocene mammals to their contemporary relatives. The biohazard associated with reviving prehistorical amoeba-infecting viruses is thus totally negligible, compared to the search for “paleoviruses” directly from permafrost-preserved remains of mammoths, woolly rhinoceros, or prehistoric horses, as it is now pursued in the Vector laboratory (Novosibirsk, Russia) [68], fortunately a BSL4 facility. Without the need of embarking on such a risky project, we believe our results with *Acanthamoeba*-infecting viruses can be extrapolated to many other DNA viruses capable of infecting humans or animals. It is thus likely that ancient permafrost (eventually much older than 50,000 years, our limit solely dictated by the validity range of radiocarbon dating) will release these unknown viruses upon thawing. How long these viruses could remain infectious once exposed to outdoor conditions (UV light, oxygen, heat), and how likely they will be to encounter and infect a suitable host in the interval, is yet impossible to estimate. But the risk is bound to increase in the context of global warming when permafrost thawing will keep accelerating, and more people will be populating the Arctic in the wake of industrial ventures.

## Author Contributions

Conceptualization and funding, C.A. and J.-M.C..; resources (samples), A.E.G., G.G, J.S., A.N.T.; virus isolation and characterization: J.-M.A., A.L.; bioinformatic analysis, O.P., M.L., S.S., J.-M.C.; writing, J.-M.C..; All authors have read and agreed to the published version of the manuscript.

## Data Availability Statement

The newly isolated viruses described here are available for collaborative studies upon formal request through a material transfer agreement.

## Acknowledgments

We thank Dr. Maksim Cheprasov (NEFU, Yakutsk, Russia) and Dr. Vladimir Pitulko (RAS, St Petersburg, Russia) for their generous gift of ancient permafrost samples. We also thank our volunteer collaborator, Alexander Morawitz, for collecting the Kamchatka soil samples and Dr. Lyubov Shmakova (Soil Cryology Lab, Pushchino, Russia) for providing samples from which, unfortunately, no viruses were rescued in the present study. We acknowledge the computing support of the PACA BioInfo platform. We thank M. Ulrich (DFG project #UL426/1-1) and P. Konstantinov for helping with fieldwork at the Yukechi site, as well as the Alfred Wegener Institute and Melnikov Permafrost Institute logistics for field support and sample acquisition.

